# BioMaster: Multi-agent System for Automated Bioinformatics Analysis Workflow

**DOI:** 10.1101/2025.01.23.634608

**Authors:** Houcheng Su, Weicai Long, Yanlin Zhang

## Abstract

**Motivation:** The rapid expansion of biological data has significantly increased the complexity of bioinformatics workflows, which often involve intricate, multi-step processes. These tasks demand considerable manual effort from bioinformaticians, creating inefficiencies and limiting scalability. Recent advancements in large language model (LLM)-powered agents offer promising solutions to streamline and automate these workflows. However, existing automated systems, while effective for short, well-defined tasks, often struggle with long, multi-step workflows due to challenges such as error propagation, limited adaptability to emerging tools, and the inability of LLMs to generalize to niche bioinformatics tasks. Achieving effective workflow automation requires robust task coordination, dynamic knowledge retrieval, and mechanisms to ensure errors are identified and resolved before they impact downstream processes.

**Results:** We present BioMaster, a multi-agent framework designed to automate and streamline complex bioinformatics workflows. BioMaster incorporates specialized agents with role-based responsibilities, enabling precise task decomposition, execution, and validation. It leverages Retrieval-Augmented Generation (RAG) to dynamically retrieve domain-specific knowledge, improving adaptability to new tools and niche analyses. BioMaster also introduces enhanced control over input and output validation to ensure pipeline consistency and employs a memory management strategy optimized for handling long workflows. Experiments across diverse bioinformatics tasks, including RNA-seq, ChIP-seq, single-cell analysis, and Hi-C processing, demonstrate that BioMaster significantly outperforms existing methods in accuracy, efficiency, and scalability. By addressing key limitations in workflow automation, BioMaster offers a robust solution for modern bioinformatics challenges.

**Availability:** https://github.com/ai4nucleome/BioMaster

## Introduction

Bioinformatics is a multidisciplinary field that applies data science techniques to interpret and analyze complex biological data, aiming to uncover biological insights and deepen our understanding of life processes Gauthier et al. (2019); Lesk (2019). Advances in high-throughput biological technologies Rhoads and Au (2015); Reuter et al. (2015); Jain et al. (2016), such as single-cell sequencing Nawy (2014), spatiotemporal omicsSchlotter et al. (2018), ATAC-seq Buenrostro et al. (2015), and 3D genome experimentsBelton et al. (2012), have transformed the scale and complexity of data generated in life sciences Reuter et al. (2015). These methods produce vast and intricate datasets that demand sophisticated analysis workflows requiring a blend of computational expertise and biological understanding Wratten et al. (2021).

However, traditional bioinformatics tools and pipelines often rely heavily on manual effort, posing challenges in terms of efficiency, reproducibility, and scalability. As the complexity of these workflows increases with the adoption of emerging technologies Nawy (2014); Subramanian et al. (2020); Belton et al. (2012); Schlotter et al. (2018); Belton et al. (2012); Schlotter et al. (2018); Ng and Kirkness (2010); Retterer et al. (2016); Ozsolak and Milos (2011); Kolodziejczyk et al. (2015); Buenrostro et al. (2015); Park (2009), the need for specialized knowledge and extensive experience has become a significant barrier. While many research institutions employ dedicated bioinformatics staff to support data analysis, it often takes years of hands-on training for students and early-career researchers to become proficient in these workflows. This steep learning curve not only slows the pace of discovery but also highlights the urgent need for automated and intelligent systems that can streamline bioinformatics analysis, reduce technical barriers, and enable researchers to focus on generating novel insights.

Recent advancements in artificial intelligence (AI), particularly the development of Large Language Models (LLMs) Zhao et al. (2023); Park et al. (2023), have opened new possibilities for automating bioinformatics workflows. Systems such as ChemCrow M. Bran et al. (2024), CellAgent Xiao et al. (2024), and AutoBA Zhou et al. (2024) exemplify the potential of AI-driven agents to tackle specialized tasks across diverse domains, including chemical synthesis, single-cell RNA sequencing (scRNA-seq) analysis, and multi-omics data processing. For instance, ChemCrow autonomously proposes production pathways for compounds, evaluates reagent prices, assesses potential hazards (e.g., toxicity or explosions), operates virtual experimental platforms, and synthesizes and analyzes chemical products M. Bran et al. (2024). Similarly, CellAgent adopts a multi-agent architecture tailored to scRNA-seq analyses, leveraging role-specific agents for task planning, execution, and validation. AutoBA, on the other hand, provides end-to-end automation for multi-omics workflows, enhancing adaptability to new tools while reducing reliance on manual intervention.

While these systems mark significant progress, challenges remain in scaling AI frameworks to manage the complexity and breadth of real-world bioinformatics workflowsHuang et al. (2024). Early systems like AutoBA relied on a single-agent approach, which, while effective for simpler workflows, struggles to meet the demands of bioinformatics tasks that require expertise across multiple domains. More recently, multi-agent systems such as CellAgent have introduced role-specific agents to better handle complex tasks, particularly for scRNA-seq data analysis. These systems utilize memory mechanisms to manage intermediate results and coordinate across steps. However, as workflows scale to include dozens of steps, memory systems can become excessively large, leading to performance degradation in LLMs, which struggle to handle overly long contexts Liu et al. (2024). Additionally, while some frameworks have incorporated Retrieval-Augmented Generation (RAG) Lewis et al. (2020) to provide domain-specific knowledge, challenges remain in aligning retrieved information with practical execution, especially in highly intricate workflows.

Despite these advancements, existing systems often fall short in managing workflows holistically, efficiently scaling to longer pipelines, and ensuring reliable execution across diverse bioinformatics tasks. Addressing these limitations is critical to advancing automation in computational biology.

To tackle these challenges, we propose BioMaster, a multi-agent framework designed to automate and scale bioinformatics workflows effectively. BioMaster integrates advanced reasoning capabilities, efficient memory control, and robust RAG mechanisms to address the limitations of existing tools. It ensures scalability, adaptability, and accuracy even for the most complex workflows, significantly reducing reliance on human intervention.

In this study, we evaluate BioMaster across a range of bioinformatics tasks and pipelines, demonstrating its ability to streamline complex workflows and reduce manual effort. By addressing the limitations of current frameworks, BioMaster takes an important step forward in advancing automation in computational biology, enabling researchers to focus more on generating meaningful biological insights.

## Materials and methods

### Overall Framework

The overall framework of BioMaster is illustrated in Fig. 1. Users start by specifying their bioinformatics analysis goal and providing the necessary input files for processing. From there, BioMaster handles the entire workflow autonomously, leveraging a multi-agent architecture to manage complex bioinformatics tasks. The process begins with the Plan Agent, which takes the user-defined goal, embeds it into a word vector, and performs similarity retrieval using the Plan RAG (Retrieval-Augmented Generation). This retrieval step supplements the goal by incorporating two highly relevant knowledge entries from the RAG database, bridging gaps in the LLM’s understanding of specialized or recent bioinformatics knowledge. The Plan Agent then decomposes the goal into a series of indivisible steps, each responsible for processing the input data and producing intermediate results, leading to the final analysis output.

**Fig. 1:**
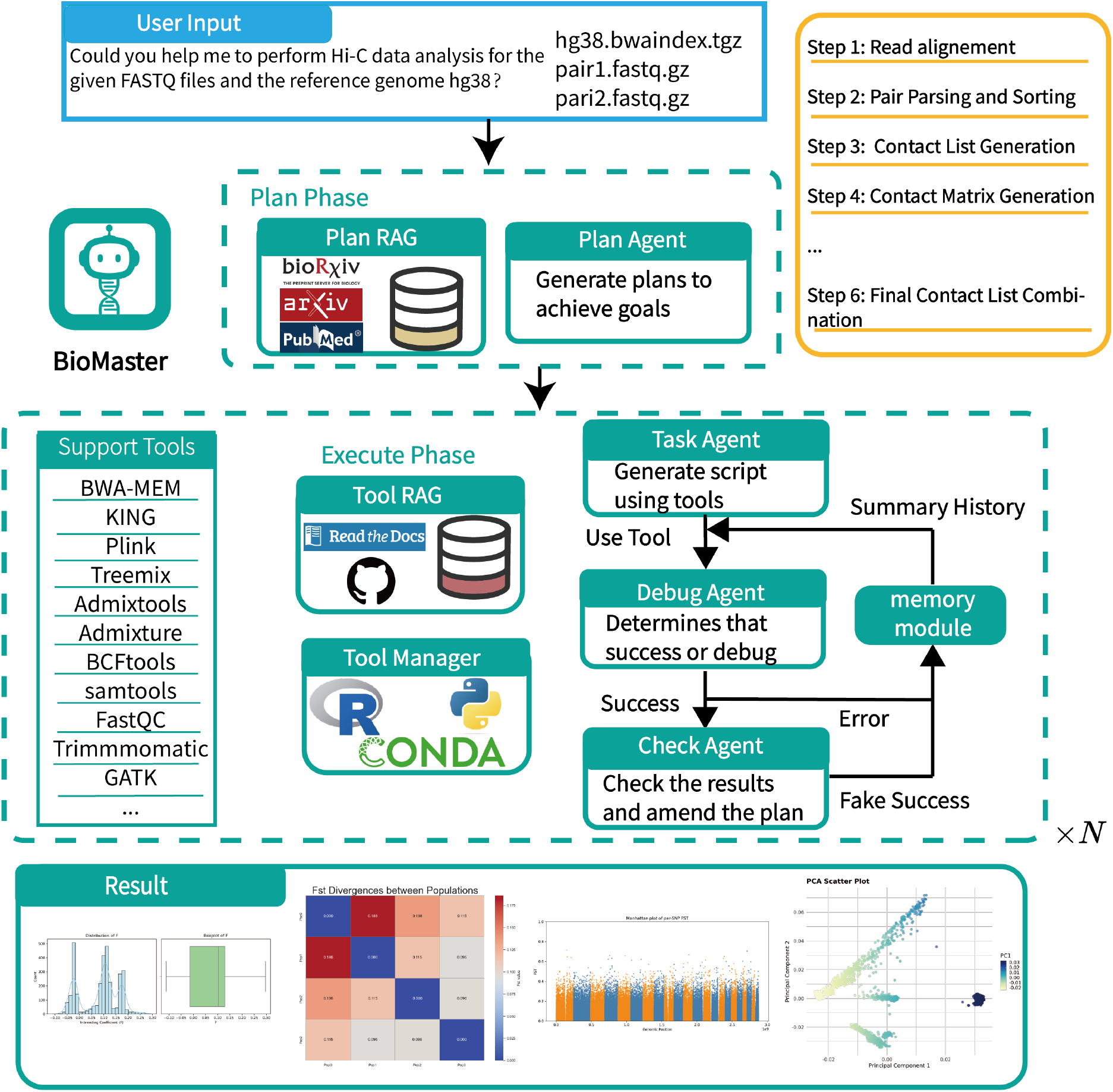
Overview of BioMaster. The workflow initiates with user-provided input, which is processed by the Plan Agent leveraging Plan RAG to design an optimized workflow. The Task Agent subsequently generates and executes scripts informed by Tool RAG knowledge. Upon completion, the Debug Agent evaluates the execution success and performs debugging as needed. The process concludes with the Check Agent validating the outputs and resolving any inconsistencies to ensure accuracy and reliability.

Once the steps are defined, the Task Agent executes the first step by integrating task descriptions from the Plan, tool requirements, input file paths, and expected output file names. The agent queries the Execute RAG to retrieve detailed descriptions and usage examples for the specified tools, enhancing its understanding of the tools’ functionality. Using this information, the Task Agent generates an executable script, including commands for tool installation and execution.

After execution, the script’s output is sent to the Debug Agent for evaluation. The Debug Agent uses information from the Plan, the Task Agent, and Execute RAG to assess whether the step was executed successfully. If errors are detected, the Debug Agent analyzes error messages and generates a new script with revised instructions. A memory module is incorporated to summarize past interactions, avoiding redundancy and conserving token usage during these iterative corrections. This process continues until the step is executed successfully.

Once the Debug Agent verifies the output, the results are passed to the Check Agent. The Check Agent ensures that the output files meet the expectations outlined in the Plan. It checks that file names match the expected naming convention, verifies that the files are non-empty, and confirms that the analysis step was executed properly. If discrepancies are found—such as mismatched file names, empty outputs, or deviations from the planned output format—the Check Agent corrects these issues and adjusts subsequent steps to prevent cumulative errors. This ensures that false completions and cascading failures are avoided throughout the workflow.

After completing the current step, the system advances to the next step in the Plan and repeats the process, cycling through the Task Agent, Debug Agent, and Check Agent. This iterative framework continues until all steps are executed successfully, resulting in the final analysis output that aligns with the user’s specified goal.

### Environment

We used OpenAI’s GPT-4 as LLMs for all agents. The system was deployed on both local and cloud servers to evaluate its performance in different computational environments. The local server specifications include an AMD EPYC 7K62 48-Core Processor and 8GB of RAM. The cloud server specifications feature a 13th Gen Intel Core i9-13900K CPU and 94GB of RAM. Since the bioinformatics workflows in our system do not heavily depend on GPU resources, no specific GPU requirements are necessary unless deploying LLaMA-based models, which require GPU devices. For optimal performance, we recommend a CPU with at least 16GB of RAM to support efficient execution of bioinformatics analyses.

The system is built on Ubuntu 22.04, with Conda serving as the primary package and tool management platform. A Python environment (version 3.12) and an R environment (version 4.4.1) were configured using Conda to ensure compatibility with bioinformatics tools and libraries. BioMaster is developed using the LangChain framework, which provides robust support for integrating and managing multi-agent workflows.

### Multi-Agent Framework

Our multi-agent framework is built using a role-based approach that defines the responsibilities and behavior of each agent through explicit instructions to the large language model. Each agent is instantiated with a specific role and operates autonomously, equipped with its own local knowledge base and local memory. Local knowledge provides task-specific information tailored to the agent’s role, while local memory tracks the agent’s historical actions, enabling efficient task management and refinement over time. To ensure modularity, the local knowledge and memory of each agent are not shared. However, all agents access a global knowledge repository containing project-wide information, such as workflow context, input files, and expected outputs, ensuring coherent collaboration across the system.

The multi-agent system integrates specialized agents to perform key functions within the bioinformatics workflow. These agents collaboratively handle the stages of planning, execution, debugging, and validation. The Plan RAG supports the planning phase by enabling agents to retrieve domain-specific workflow knowledge, addressing knowledge gaps in the LLM for specialized or newer bioinformatics tools. The Execute RAG enhances task execution and debugging by providing tool-specific guidance and examples.

Collaboration between agents occurs within a shared environment, where agents exchange information while maintaining role-specific autonomy. For example, the planning process defines a sequence of tasks that is executed and refined step-by-step, with each agent performing its role in the workflow and sharing results for subsequent steps. The framework is designed to ensure modularity and adaptability, allowing it to handle complex bioinformatics tasks with minimal overlap or redundancy.

This design, leveraging role-based agents, retrieval-augmented generation, and collaborative task execution, enables BioMaster to effectively manage diverse and multi-step workflows while maintaining scalability and robustness.

### Retrieval-Augmented Generation (RAG)

In BioMaster, we implemented RAG to enhance the system’s ability to handle bioinformatics workflows by retrieving domain-specific knowledge. RAG is integrated into both the planning and execution phases of the framework, enabling the agents to access relevant information and make informed decisions.

During the planning phase, the Plan RAG module is used by the Plan Agent to retrieve workflow-related knowledge from curated databases. This includes detailed descriptions of bioinformatics methods, best practices, and up-to-date tools. By supplementing the user-defined goal with this retrieved knowledge, the Plan Agent ensures the generated workflows are both accurate and aligned with the latest advancements in bioinformatics.

For the execution phase, we designed the Execute RAG module, which is used by the Task Agent and Debug Agent. Execute RAG provides tool-specific information, including parameter descriptions, usage examples, and error-handling strategies. This detailed guidance enables the Task Agent to generate precise commands and the Debug Agent to resolve errors effectively. Both RAG modules are configured to return the two most relevant knowledge entries for any given query, ensuring a balance between precision and computational efficiency.

To support RAG, we developed a knowledge database that includes curated content from bioinformatics tool documentation, research articles, and workflow repositories. The database is structured to enable fast and context-aware retrieval, ensuring agents can quickly access the information they need for each step of the workflow.

Our RAG system, supported by curated and user-extendable content, ensures BioMaster can flexibly adapt to new tools and methods while maintaining consistent performance across diverse bioinformatics tasks.

### Plan Agent

The Plan Agent in BioMaster is designed to generate structured workflows by decomposing user-defined goals into actionable steps. To achieve this, the agent uses a Retrieval-Augmented Generation (RAG) model, named Plan RAG, to access domain-specific bioinformatics knowledge (Fig. 2) and supplement the LLM’s understanding of the task.

**Fig. 2:**
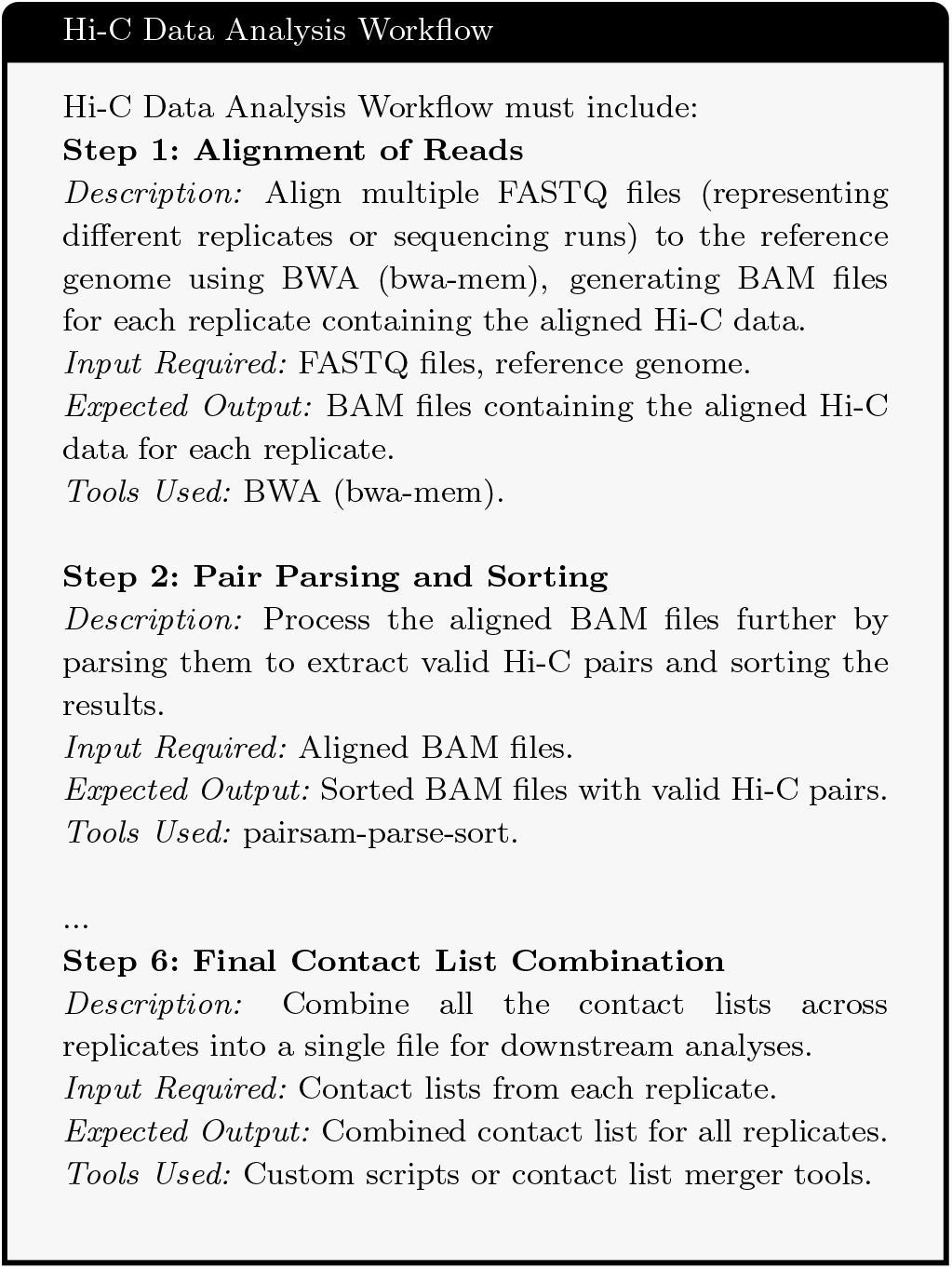
An example record included in the knowledge database of our plan RAG.

When a user provides an input goal, the text is processed using OpenAI’s text-embedding-3-large model to generate embedding vectors. These vectors are then compared with those stored in the Plan RAG database using cosine similarity. The similarity computation identifies the top two most relevant entries in the database, which are retrieved to support the planning process. The Plan RAG database contains curated knowledge, including standardized workflows, tool documentation, and supplementary materials. To maintain adaptability, the system allows users to add new knowledge sources, such as PDFs, which are automatically converted to text, summarized by the LLM, and stored for future retrieval.

The Plan Agent integrates the retrieved knowledge with the user input to create a detailed workflow plan (Fig. 3). The plan defines each task in the workflow, specifying the tools required, expected input files with descriptions and paths, and the expected outputs. The final plan is formatted in JSON to ensure compatibility with downstream agents for execution and debugging. This structured output reduces ambiguity and ensures the workflow can be executed seamlessly.

**Fig. 3:**
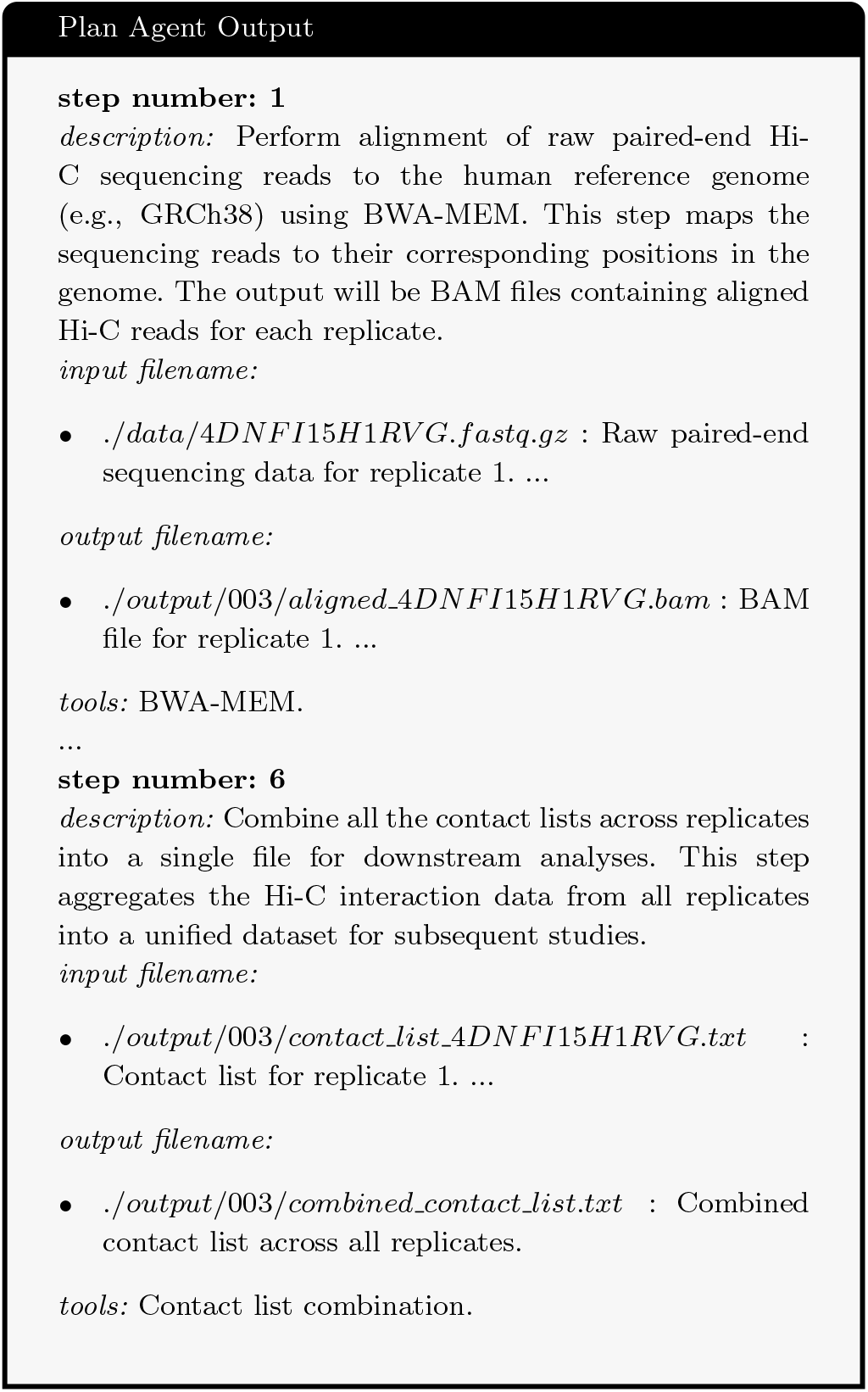
An example plan produced by the plan agent.

The Plan Agent’s reliance on Plan RAG ensures that it can incorporate up-to-date methods and tools, adapting to changes in the bioinformatics landscape. This dynamic retrieval process, combined with the clear and structured representation of the workflow, enables BioMaster to handle complex bioinformatics tasks with greater reliability and flexibility.

### Task Agent and Debug Agent

The Task Agent and Debug Agent play complementary roles in the execution phase of BioMaster. The Task Agent is responsible for translating the steps defined by the Plan Agent into executable commands, while the Debug Agent ensures these commands are executed successfully by identifying and resolving errors.

When the Plan Agent decomposes the user’s goal into steps, each step is passed sequentially to the Task Agent. The Task Agent retrieves relevant knowledge from the Execute RAG, which contains detailed tool descriptions, parameter guidance, and usage examples. Using this retrieved information and the provided step description, the Task Agent generates an executable script tailored to the task. This process ensures that the commands account for both tool-specific requirements and the broader workflow context. Once the script is generated, the Task Agent executes the commands and sends the outputs, along with the script and related input details, to the Debug Agent for validation. The Debug Agent verifies whether the execution was successful. If errors occur, it retrieves troubleshooting information from the Execute RAG, analyzes the error messages, and modifies the script to address the issue. This iterative process continues until the execution succeeds. By leveraging role-specific retrieval and its ability to refine scripts dynamically, the Debug Agent minimizes failures and ensures that the workflow progresses without interruptions.

To optimize memory usage during iterative execution, we implemented memory summarization. Instead of retaining all tokens from prior steps, which would lead to inefficiencies and redundancies, the Debug Agent summarizes key points from previous interactions using LLM-based techniques. This allows the system to conserve resources while preserving the essential context required for accurate debugging and script refinement.

After successful execution, the Debug Agent passes the validated output to the Check Agent for further verification. This modular and iterative collaboration between the Task Agent and Debug Agent ensures that each step of the workflow is executed with precision, reliability, and efficiency.

If successful, the output is passed to the Check Agent. The Fig. 4 shows an output example of the Debug Agent.

**Fig. 4:**
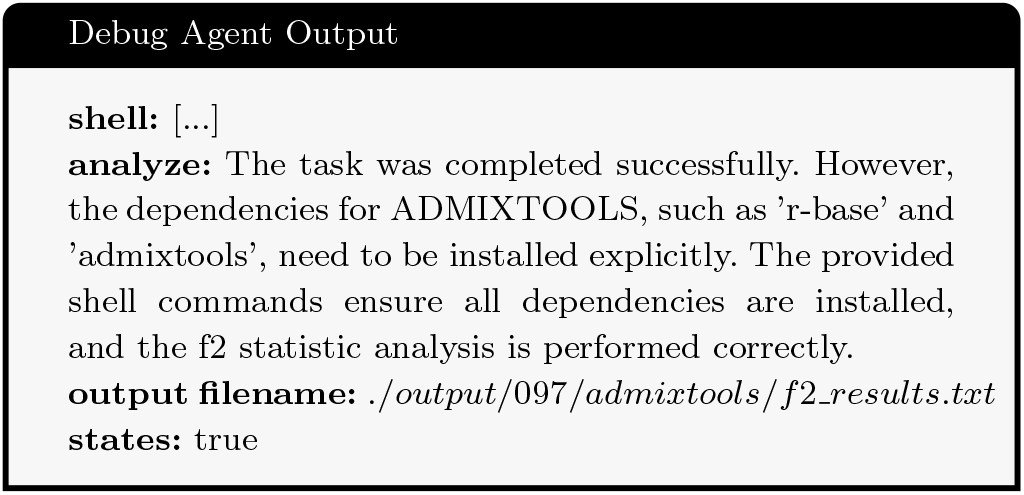
An example output from our Debug Agent.

### Check Agent

Through our testing, we found that for long tasks, there can be deviations between actual and expected results. As the number of steps increases, these deviations accumulate, eventually causing the workflow to fail. The Check Agent in BioMaster is designed to verify the reliability of outputs after each execution step, ensuring errors are identified and corrected before they propagate to subsequent steps. After each step, the Check Agent validates the output files generated by the Debug Agent against the expectations specified in the Plan. It first checks for the existence of the output files in the designated folder and ensures that the files are non-empty. If these conditions are not met, the Check Agent marks the step as unsuccessful and sends feedback to the Debug Agent for correction.

In addition to validating the Debug Agent’s outputs, the Check Agent compares the confirmed outputs with the Plan Agent’s original expectations. If discrepancies are found, such as differences in file formats, naming conventions, or missing steps, the Check Agent updates the plan to ensure compatibility with downstream tasks. This prevents small errors, such as mismatched file formats or incomplete processing steps, from affecting subsequent analysis steps.

By performing these validations after every step, the Check Agent ensures that workflows remain aligned with the original plan and that errors are addressed immediately. This iterative validation and feedback mechanism enhances the reliability of the workflow, ensuring accurate and complete execution of complex bioinformatics tasks.

## Results

### BioMaster is more accessible than alternative agents

To evaluate the performance of BioMaster, we conducted comparative experiments against AutoBA and OpenAI’s ChatGPT.

To ensure fairness, all methods utilized the same underlying large language model (LLM): the GPT-4o model (version ‘gpt-4o-2024-08-06’). Debugging procedures varied across methods. BioMaster and AutoBA employed fully automated debugging processes, whereas ChatGPT required manual intervention, with error messages copied and pasted into the chat interface for resolution. Each method was allowed up to five debug attempts per step, and tasks that were not successfully completed within this limit were recorded as failures.

The evaluation covered a diverse range of bioinformatics tasks, including ChIP-seq analysis, RNA-seq analysis, single-cell data analysis, population genetic analysis, spatial transcriptomics, and Hi-C genomic analysis. A total of 20 distinct experiments were conducted to comprehensively assess BioMaster’s robustness and versatility, the results are shown in Table 1. For ChIP-seq analysis, data from PRC1 and PRC2 members in mouse embryonic stem cells were used, with sample IDs SRR620204, SRR620205, SRR620206, and SRR620208 Morey et al. (2013). RNA-seq analysis used mouse data from the Sequence Read Archive (SRA) with sample IDs SRR1374921, SRR1374922, SRR1374923, and SRR1374924 Haas et al. (2019). Single-cell analysis employed publicly available Peripheral Blood Mononuclear Cells (PBMC) data from 10X Genomics. Population genetic analysis utilized data from the 1000 Genomes Project Consortium et al. (2012), focusing on the YRI, CEU, and CHB populations. Hi-C genomic analysis relied on GM12878 cell data with MboI digestion, including sample IDs 4DNFI15H1RVG, 4DNFIZHUKESO, 4DNFIKVDGNJN, and 4DNFIEQ58J6G Rao et al. (2014).

**Table 1.** Agents performance on various tasks, where ✓ indicates successful execution and *×* indicates failed execution.

| Bioinformatics Pipelines | Task | AutoBA | ChatGPT | BioMaster |
| --- | --- | --- | --- | --- |
| ChIP-seq data analysis | Peak calling | ✓ | ✓ | ✓ |
| ChIP-seq data analysis | Discover motifs within the peaks | ✓ | ✓ | ✓ |
| ChIP-seq data analysis | Functional Enrichment | × | × | × |
| RNA-seq Analysis | Find Differentially expressed genes | ✓ | ✓ | ✓ |
| RNA-seq Analysis | Identify the top5 downregulated gene | ✓ | ✓ | ✓ |
| RNA-seq Analysis | Predict Fusion genes using STAR-Fusion | ✓ | ✓ | ✓ |
| RNA-seq Analysis | RNA editing | × | ✓ | ✓ |
| Single-cell data analysis | Find the differentially expressed genes | ✓ | ✓ | ✓ |
| Single-cell data analysis | find marker genes based on count matrix | ✓ | × | × |
| Single-cell data analysis | Perform clustering | ✓ | ✓ | ✓ |
| Single-cell data analysis | Identify top5 marker genes | ✓ | ✓ | ✓ |
| Spatial transcriptomics | Neighborhood enrichment analysis | ✓ | ✓ | ✓ |
| Population genetic analysis | Data control and statistics | ✓ | ✓ | ✓ |
| Population genetic analysis | Smartpca Analysis | × | × | ✓ |
| Population genetic analysis | Runs of Homozygosity (ROH) | × | ✓ | ✓ |
| Population genetic analysis | Treemix Analysis | × | × | ✓ |
| Hi-c genetic analysis | Hi-C data mapping, sorting and conversion | ✓ | ✓ | ✓ |
| Hi-c genetic analysis | Hi-C Data Pair Parsing and Cleaning | × | × | ✓ |
| Hi-c genetic analysis | Hi-C Data Contact Matrix Creation and Processing | ✓ | ✓ | ✓ |
| Hi-c genetic analysis | Hi-c Fully preprocessing process | × | × | ✓ |

Mainstream bioinformatics workflows, such as RNA-seq, ChIP-seq, single-cell analysis, and spatial transcriptomics, generally involve around five steps. Across these tasks, BioMaster, AutoBA, and ChatGPT performed comparably well, with failures typically resulting from Conda dependency conflicts, which required environment adjustments—a challenge even for human experts. However, as the complexity of workflows increased, performance gaps between the methods became apparent.

For population genetics tasks Harney et al. (2021), such as executing smartPCA and treemix Pickrell and Pritchard (2012), workflows required extensive preprocessing, including data cleaning and parental removal, as well as correctly formatting parameter files. Errors in handling these niche tasks frequently arose due to missing steps or file mismatches, highlighting limitations in existing LLM-based systems for complex and domain-specific analyses.

Hi-C data preprocessing, involving at least ten steps, presented further challenges. While all methods successfully managed basic tasks such as mapping, sorting, and contact matrix creation, failures occurred during advanced steps, including pair parsing and file cleaning. Fig. 5 illustrates two key observations. In Fig. 5(a), both AutoBA and ChatGPT struggled with file merging due to incomplete planning. Even when provided with the official 4DN pipeline, they encountered errors because of insufficient detail for intricate workflows. We then studied how each agent performed in generating commands. To make a fair comparison, we provided all agents with expert-edited plans (i.e, all plans are correct). In Fig. 5(b), ChatGPT, despite leveraging correct plan, struggled with file path handling, while AutoBA failed to verify outputs, resulting in cascading errors. BioMaster’s Check Agent addressed these issues by validating outputs at each step, preventing error propagation, and ensuring task completion.

**Fig. 5:**
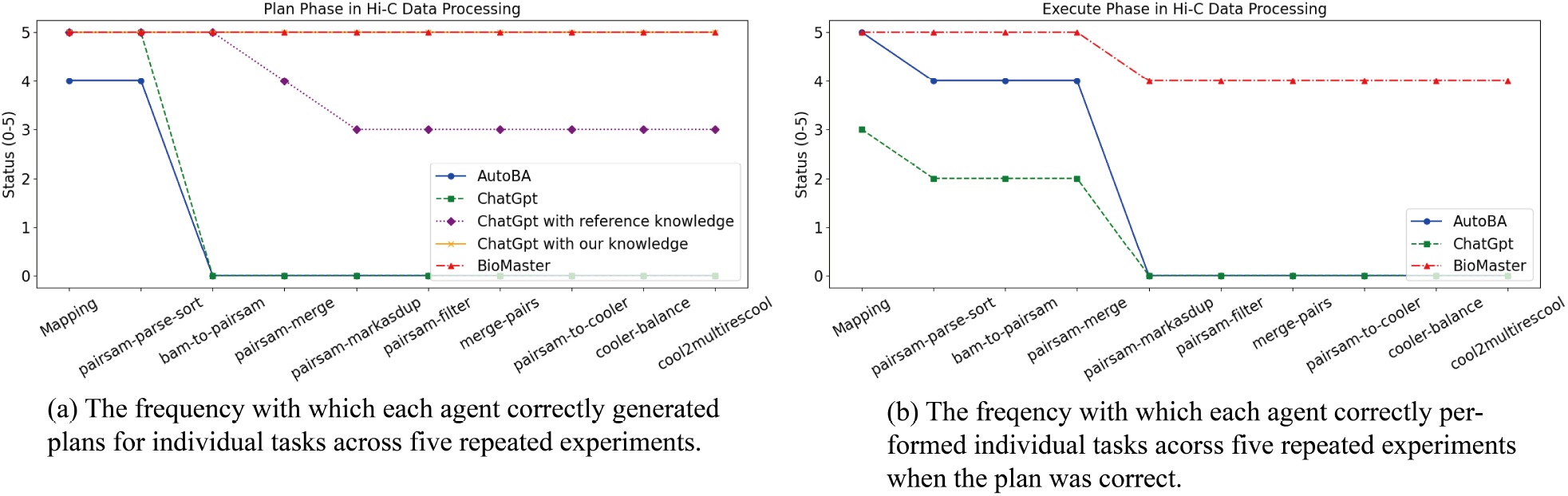
A comparison of various agents on performing a typical Hi-C data analysis workflow from the 4DN project Reiff et al. (2022). Reference knowledge refers to the original workflow provided by 4DN, while our knowledge represents the insights and workflows created by BioMaster, which were derived by incorporating 4DN’s workflow as input.

These results highlight BioMaster’s ability to handle complex, multi-step workflows more effectively than alternative methods. Its modular design and integration of expert knowledge through mechanisms like Plan RAG and the Check Agent enable it to address challenges that other systems cannot, providing a robust solution for bioinformatics analysis. The findings underscore the importance of integrating domain expertise and rigorous validation mechanisms in building reliable bioinformatics agents.

### Ablation experiment

In this section, we present an ablation study to assess the contribution of key components in our model, specifically the Plan RAG, Tool RAG, and Check Agent, to the overall performance.

For Plan RAG and Check Agent, the primary objective is to improve the accuracy and reliability of long workflows while minimizing discrepancies between planned and executed steps. To evaluate their impact, we conducted an ablation study by systematically removing these modules during Hi-C preprocessing tasks. The results, illustrated in Fig. 6(a), reveal that the absence of Plan RAG significantly hampers the system’s ability to generate effective and detailed plans, leading to incomplete or suboptimal workflows. Similarly, the removal of the Check Agent has minimal impact on shorter tasks but severely affects the success of longer workflows, particularly when small deviations occur in intermediate outputs. These findings underscore the critical role of Plan RAG in effective workflow planning and the Check Agent in ensuring consistency and preventing cascading errors in complex workflows.

**Fig. 6:**
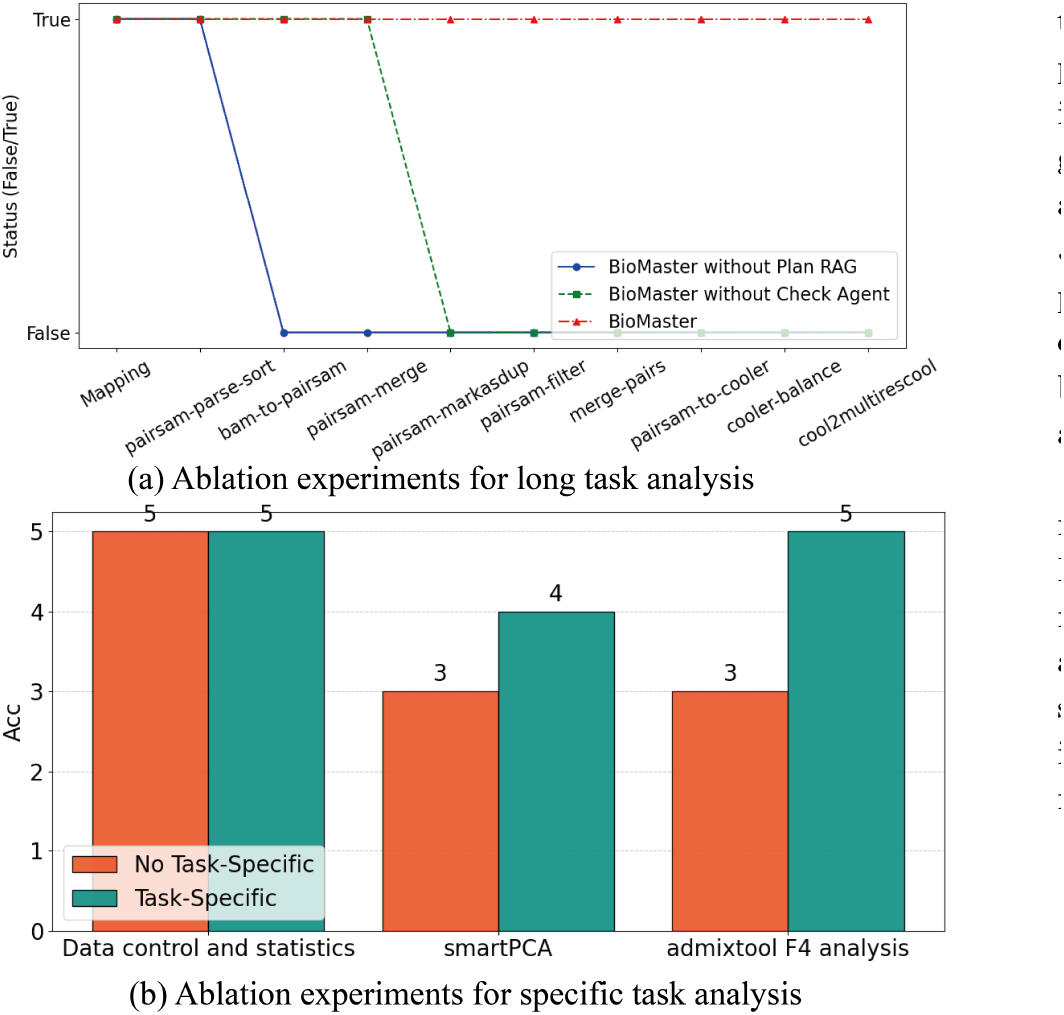
Ablation experiment result.

For Tool RAG, although the concept itself is not novel, we enhanced its functionality by incorporating detailed examples and specific instructions tailored to bioinformatics tasks. To assess its effectiveness, we conducted an ablation study by removing these enhancements and evaluated the model on population genetics tasks such as smartPCA Patterson et al. (2006) and admixtools F4 analysis Maier et al. (2023). As shown in Fig. 6(b), the absence of these Tool RAG enhancements had minimal impact on simpler tasks. However, in more complex analyses like smartPCA and admixtools F4, the performance significantly deteriorated, validating the importance of the enhanced Tool RAG in improving accuracy and reliability in intricate bioinformatics tasks.

These results collectively demonstrate that the Plan RAG, Tool RAG, and Check Agent are essential components of our framework, each addressing distinct challenges in planning, execution, and error handling. Their integration enables BioMaster to excel in managing both simple and complex workflows, ensuring robust and accurate bioinformatics analysis.

## Discussion and Conclusion

In this work, we explored novel approaches to developing bioinformatics agents by introducing BioMaster, a multi-agent framework designed to address the complexity of modern bioinformatics workflows. Our experimental evaluation demonstrates that BioMaster outperforms existing frameworks across a range of bioinformatics tasks, achieving higher success rates in executing complex workflows. Key contributions of this work include improved handling of inputs and outputs, the development of a memory mechanism tailored to workflow management, drawing inspiration from the mem0 memory model to enhance memory control in complex processes, and the integration of specialized agents to address different stages of analysis.

BioMaster’s modular, multi-agent design provides notable advantages over single-agent frameworks. By assigning distinct roles to each agent and tailoring their capabilities—such as dynamic task planning, error detection and correction, and output validation—the system achieves better task decomposition and execution reliability. Additionally, the incorporation of retrieval-augmented generation (RAG) significantly enhances the system’s ability to acquire and apply domain-specific knowledge. We emphasize the need to design different types of RAG modules and deliver varying levels of detail to individual agents, ensuring that each agent performs its role effectively while contributing to the overall workflow.

Comparative and ablation studies validate the effectiveness of BioMaster’s components, such as the Plan RAG module, which improves task planning, and the Check Agent, which ensures the reliability of intermediate outputs. These innovations collectively enable the system to mitigate common issues in long workflows, such as error accumulation and inefficiencies, while enhancing both efficiency and accuracy.

This work highlights promising directions for future development in bioinformatics agents. While BioMaster demonstrates the benefits of multi-agent frameworks and RAG-enhanced workflows, further research is needed to refine memory control, optimize agent collaboration, and expand the framework’s adaptability to new tools and datasets. By addressing these challenges, the principles explored in this study can help pave the way for more sophisticated and scalable AI-driven bioinformatics systems.

## Author contributions

H.S. and Y.Z. conceived the study. H.S. and W.L. performed the analysis. Y.Z. supervised the project. H.S. and Y.Z. wrote the article. All authors read and approved the final article.

